# Moonlighting adenylyl cyclases in plants – an *Arabidopsis thaliana* 9-*cis*-epoxycarotenoid dioxygenase as point in case

**DOI:** 10.1101/2021.03.23.436544

**Authors:** Inas Al-Younis, Aloysius Wong, Basem Moosa, Mateusz Kwiatkowski, Krzysztof Jaworski, Chris Gehring

## Abstract

Adenylyl cyclases (ACs) and their catalytic product cAMP are regulatory components of plant responses. AC domains are intrinsic components of complex molecules with multiple functions, some of which are co-regulated by cAMP. Here we used an amino acid search motif based on annotated ACs in organisms across species to identify 12 unique *Arabidopsis thaliana* candidate ACs, four of which have a role in the biosynthesis of the stress hormone abscisic acid (ABA). One of these, the 9-*cis*-epoxycarotenoid dioxygenase (NCED3, At3g14440), was identified by sequence and structural analysis as a putative AC and then tested experimentally for activity. We show that an NCED3 AC fragment can complement an AC deficient *E. coli* mutant and this rescue is nullified when key amino acids in the AC motif are mutated. AC activity was also confirmed by tandem liquid chromatography mass spectrometry (LC-MS/MS). Our results are consistent with a moonlighting role for mononucleotide cyclases in multi-domain proteins that have at least one other distinct molecular function such as catalysis or ion channel activation and promise to yield new insights into tuning mechanisms of ABA dependent plant responses. Finally, our search method can also be applied to discover ACs in other species including *Homo sapiens*.

**Highlights:**

- An adenylyl cyclase (AC) catalytic center motif identifies novel ACs in plants
- ACs can moonlight in complex proteins with other enzymatic domains
- A 9-*cis*-epoxycarotenoid dioxygenase essential for abscisic acid synthesis contains an AC
- This finding implicates cAMP in abscisic acid synthesis and signaling

## 1. Introduction

In plants, cyclic nucleotide monophosphates and their cyclases are gaining increasing attention due to their involvement in crucial developmental and physiological processes including regulating guard cell movements and responses to abiotic and biotic stresses [1]. Furthermore, the *Arabidopsis thaliana* cyclic nucleotide interactome harbors proteins that cross-talk with cyclic nucleotides, nitric oxide (NO) and other reactive oxygen species to signal for plant defense responses [2].

Given the possibility that many ACs in *Arabidopsis thaliana* remain undetected, we have used a novel motif-based approach to identify candidate ACs [3]. This motif is based on conserved amino acids in catalytic centers of proteins across species [4] and has since identified numerous nucleotide cyclases (for review see: [1]). The same method has also been applied to discover functional NO binding [5–8] and ABA binding sites [9] in *Arabidopsis* proteins.

Here we built a stringent AC catalytic center motif, queried the *Arabidopsis thaliana* proteome and identified a number of candidate ACs of which one, a 9-*cis*-epoxycarotenoid dioxygenase (NCED3, At3g14440), was selected for computational and functional analysis since NCEDs are crucial components of ABA biosynthesis [10] and hence, also plant stress responses. Finally, we propose a model that can guide further investigations of the role of cAMP in ABA-dependent responses.

## 2. Materials and Methods

### 2.1 Structural analysis of AtNCED3 AC and NCED3 domains

AtNCED3 was modeled against the crystal structure of a maize viviparous14 protein (PDB ID: 3NPE) using MODELLER (ver. 9.25) [11] and docking simulations of ATP to the AC center of AtNCED3 was performed using AutoDock Vina (ver. 1.1.2) [12]. In docking simulations, all bonds in the ATP were allowed to move freely but the protein was set rigid. Docking orientations of ATP were evaluated based on a previously ascertained “correct binding pose” where adenine points towards position 1, which normally occupies the interior of the AC pocket, and phosphate points towards position 14, which normally occupies the solvent-exposed entrance area of the AC pocket [3]. Docking simulations consider both spatial and charge in the vicinity of the catalytic center based on pre-determined grids that cover the entire AC center and can afford free rotation of ATP substrate which we have set prior to docking experiments. The ATP docking poses were analyzed, and all images created by UCSF Chimera (ver. 1.10.1) [12]. Chimera was developed by the Resource for Biocomputing, Visualization, and Informatics at the University of California, San Francisco (supported by NIGMS P41-GM103311).

### 2.2 Complementation of an AC deficient E. coli mutant

pDEST17-AtNCED3^211-440^ constructs (see Supplementary Methods) were transformed into the *E. coli* cyaA mutant SP850 strain [lam-, el4-, relA1, spoT1, cyaA1400 (:kan),thi-1] [13] deficient in its AC (cyaA) gene. (*E. coli* Genetic Stock Center, Yale University, New Haven, CT, US, Accession Number 7200). The transformation with the plasmid was done by heat shock (2 minutes at 42°C). Bacteria with *E. coli* cyaA mutant strain re-grown at 37°C in Luria Broth media supplemented with 100 μg/mL ampicillin and 100 μg/mL kanamycin until they reached an OD_600_ of 0.6 and then incubated with 0.5 mM isopropyl-β-D-1-thiogalactopyranoside (IPTG) for 4 hours for transgene induction prior to streaking on MacConkey agar.

### 2.3 AC biochemical assay and mass spectroscopic detection of cAMP

Cyclic AMP was generated from reaction mixture containing 10 μg of purified recombinant protein AtNCED3^211-440^ or the mutated protein (AtNCED3^S311P/D313T^), 50 mM Tris–HCl (pH 8.0), 2 mM isobutylxanthine (IBMX; Sigma-Aldrich, St. Louis, MO, US), 5 mM MnCl_2_ and 1 mM ATP at room temperature and in a final volume of 100 μL. The reaction was stopped by adding 10 μL of ≥ 4 mM EDTA. The cAMP produced was measured with the Biotrack enzyme immunoassay using the acetylation protocol as described by the manufacturer (GE Healthcare, Little Chalfont, UK). Cyclic AMP was also detected using liquid chromatography tandem mass spectrometry (LC-MS/MS) on an LTQ Orbitrap Velos mass spectrometer (Thermo Fisher Scientific, Waltham, MA, USA) [14,15]. Briefly, separation was achieved by a Sepax SFC Cyano column (150 × 4.6 mm × 5 μm) at ambient temperature, an isocratic mix of 10 mM ammonium acetate and acetonitrile (HPLC-MS grade, ratio: 60%/40%) and for the detection, positive ESI as ionization was used. Quantitation was based on the chromatographic peak areas of the samples using the extracted ion chromatogram of the product ion m/z 330.12 [15]. All enzymatic reactions were done alongside no-protein controls and they were also subjected to LC-MS/MS detections. Signal contributions by the controls are considered as “background” which we have subtracted to obtain the actual cAMP amounts.

## 3. Results and Discussion

### 3.1 Identification of candidate ACs

We assembled a 14-amino acid long AC catalytic center motif containing the key amino acids that are conserved and have direct roles in substrate binding and/or catalysis (Fig. 1A). These amino acids are separated by gaps as determined in the alignment of catalytic centers of annotated ACs in diverse prokaryotic and eukaryotic organisms. Notably, this AC search motif and its derivatives have successfully identified similar and experimentally confirmed ACs [16,17].

**Fig. 1.**
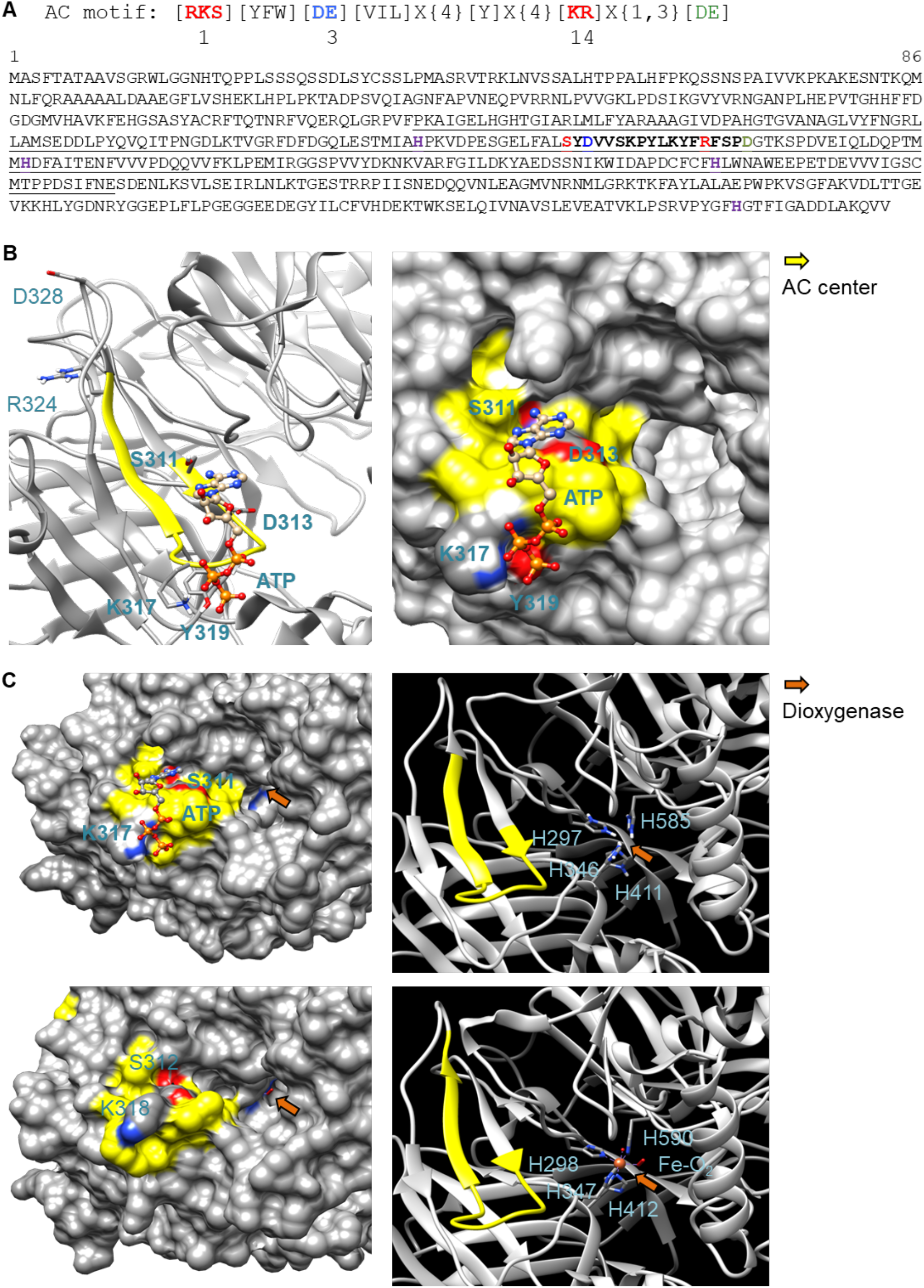
Sequence and structural analysis of 9-*cis*-epoxycarotenoid dioxygenase AtNCED3. (**A**) AC search motif and amino acid sequence of AtNCED3. The AC center motif (S311 – R324) is bolded in the amino acid sequence where position 1 (red) forms hydrogen bond with adenine, position 3 (blue) confers substrate specificity (D or E for ATP) and position 14 stabilizes the transition (ATP/cAMP). Amino acid for Mg^2+^/Mn^2+^ binding is labeled green. In the motif specific for ACs, the [DE] in position 3 (blue) allows for ATP binding. Amino acids in the square brackets denote amino acids allowed in this position, “X” denotes any amino acid and curly brackets ({}) denotes the number of undetermined amino acids. Underlined amino acids indicate the AtNCED3 fragment that was cloned and expressed for functional studies *in vitro*. The histidine residues marked in magenta are required for the octahedral binding of Fe^2+^ which is required for dioxygenase activity. (**B**) Ribbon and surface models of AtNCED3 AC center docked with ATP. The AC catalytic center is highlighted in yellow and key amino acids in the motif (except for D328 and R324) and amino acids that may interact with ATP are labeled, colored according to their charges, and shown as individual atoms in the ribbon models. AtNCED3 structure was modeled against the crystal structure of a maize viviparous14 protein (PDB ID: 3NPE). (**C**) Comparison of the AC and dioxygenase domains in AtNCED3 (top) and ZmNCED3 (bottom). AtNCED3 docked with ATP is represented as a surface model (left) with its AC and iron binding histidine amino acids at the dioxygenase domain represented as a ribbon model (right) in the top panel with corresponding regions in the ZmNCED3 structure shown in the bottom panel. AC center is highlighted in yellow and key amino acids in the motif are labeled in bold and colored according to their charges in the surface models. The iron binding histidine residues in the dioxygenase domain (orange arrows) are labeled in the ribbon models.

When the *Arabidopsis thaliana* proteome was queried with this search term, it returned 12 proteins (Supplementary Table S1) one of which, a clathrin assembly protein (CLAP, At1g68110), has been shown to have AC activity [18]. We also noted significant enrichments in several gene ontology terms (GO) notably “biosynthetic process” (GO:0009058; p=0.0064, FDA=0.046). We selected AtNCED3 (At3g14440) for further analysis since it has (11’,12’) 9-*cis*-epoxycarotenoid cleavage activity and catalyzes the first step of ABA biosynthesis from carotenoids and in doing so, enabling plant response to water stress [19–23]. AtNCED3 is therefore a promising candidate for research that examines the link between plant stress, ABA, and cAMP.

### 3.2 Structural analysis of AtNCED3 AC and NCED3 domains

Since the structure of AtNCED3 is yet to be determined, we built a model for AtNCED3 using homology modeling approach [24] (see section 2.1) to analyze the AC and NCED3 domains. AtNCED3 was modeled against the crystal structure of a maize NCED3 (ZmNCED3) which is a key enzyme in the biosynthesis of the phytohormone ABA (PDB ID: 3NPE). At 69% identity and covering 86% of AtNCED3 amino acid sequence, ZmNCED3 was the best template option identified using the BLASTp tool available at https://blast.ncbi.nlm.nih.gov/Blast.cgi?PAGE=Proteins [25], that compared AtNCED3 against a database of protein crystal structures in the protein databank (PDB). Based on the model, the AC center motif [RKS][YFW][DE][VIL]X{4}[Y]X(4)[KR]X{1,3}[DE] appears between S311 to R324 with the predicted cation binding amino acid D328 located four residues downstream of the catalytic center (Fig. 1A). Notably, the AC prediction tool ACPred available at http://gcpred.com/acpred/main.php [26] also identified with high confidence this same region as a candidate AC.

The secondary structure of the AC center assumes a slightly different fold compared to other experimentally validated AC centers that contain a helix-loop secondary fold (Fig. 1B) [27,28]. The AC center occupies a distinct pocket that docks ATP with a mean binding affinity of −4.64 ± 0.03 kcal/mol calculated from a total of 10 positive docking solutions predicted by AutoDock Vina (Fig. S1). ATP assumes an orientation much like in experimentally validated AC centers [17,27,28] where the adenine of ATP points towards position 1, which occupies the interior of the AC pocket, and the phosphate which points towards position 14, which occupies the solvent-exposed entrance area of the AC pocket (Fig. 1B,1C). We refer this substrate orientation as a “correct binding pose” [3]. However, spatial evaluation of its binding pose suggests that the phosphate end of ATP could interact more feasibly with K317 rather than R324 which, together with the predicted metal coordinating D328 residue, are seemingly too distant for interactions with ATP. Instead, the aromatic ring of a tyrosine (Y319) that is located two amino acids downstream of K317, could offer metal ion coordination (Fig. 1B). Since binding pose of ATP to ACs identified through this motif has been previously ascertained [17,18], we analyzed all docking solutions across two independent simulations to determine the positive binding pose frequency which in this case is 55.6%, thus lending confidence to the prediction that this protein can function as an AC. Docking clusters and data, and representative structures are presented in Supplementary Fig. S1.

AtNCED3 AC does not resemble canonical ACs since the motif only includes the key amino acids at the catalytic centers of conventional ACs. Moreover, ACs identified by this motif approach occupy moonlighting sites that are separate from primary domains that range from kinases to transporters and gas-sensing regions [8,14,27,28]. Due to these diverse structural architectures, these ACs assume structural folds that do not resemble the conventional classes of ACs which are often stand-alone proteins [26,29]. Thus, while structural evaluation supports favorable binding with ATP which is a pre-requisite for catalysis, there were also structural features that are unique to the AC center of NCED3.

### 3.3 Experimental validation of AtNCED3 AC activity

Functional analysis of AtNCED3 AC was conducted using two methods. The first is based on the rescue of an *E. coli* mutant (*cyaA*) deficient in its AC activity. AtNCED3^211-440^ was cloned and expressed in *E. coli* SP850 strain lacking the c*yaA* gene which prevents lactose fermentation. Due to the *cyaA* mutation, the AC deficient *E. coli* and the un-induced transformed *E. coli* cells remain colorless when grown on MacConkey agar (Fig. 2A). In contrast, the AtNCED3^211-440^ transformed *E. coli* SP850 cells, when induced with 0.5 mM IPTG, form red colored colonies much like the WT *E. coli*, indicating that a functional AC center in the recombinant AtNCED3^211-440^ has rescued the *E. coli cyaA* mutant. Significantly, a double mutant AtNCED3^S311P/D313T^ is unable to rescue the *E. coli cyaA* mutant (Fig. 2A). S311 in position 1 of the AC motif is predicted to hydrogen bond with ATP while D311 in position 3 is hypothesized to confer substrate specificity. This implies that AtNCED3^211-440^ contains a functional AC that can complement the *E. coli cyaA* mutant and that S311 and/or D311 is critical for catalysis of similar enzymatic centers [17,27,28].

**Fig. 2.**
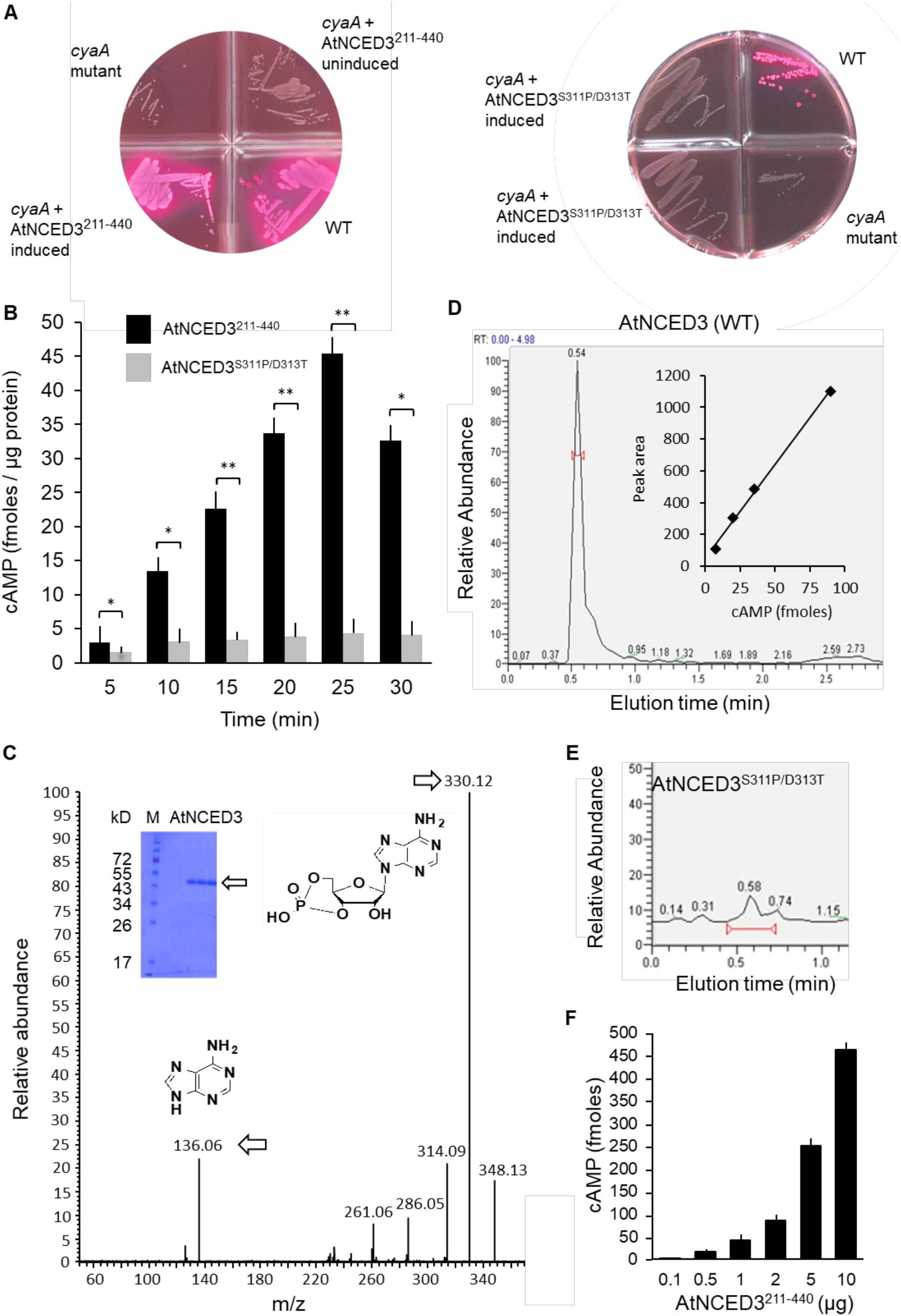
Experimental validation of AC activity *in vitro*. (**A**) Complementation of *E. coli cyaA* with AtNCED3 AC fragments AtNCED3^211-440^ (left) and AtNCED3^S311P/D313T^ (right). (**B**) cAMP generated by AtNCED3^211-440^ and AtNCED3^S311P/D313T^. Reaction mixtures contain 10 μg of AtNCED3^211-440^ or AtNCED3^S311P/D313T^, 50 mM Tris-HCl pH 8; 2 mM IBMX, 1 mM ATP and 5 mM MnCl_2_. Measurements of three independent experiments are represented as mean ± SE and were subjected to unpaired, one-tailed Student’s *t*-test (Two-Sample Assuming Unequal Variances) where * = *P* < 0.05 and ** = *P* < 0.005 compared to the activities of AtNCED^211-4403^ at the corresponding time points. Measurements at each time point compared to that at the first time point (5 minutes) for AtNCED3^211-440^ and AtNCED3^S311P/D313T^ are also statistically significant (*P*<0.05; *n*=3). (**C**) Mass spectrometric detection of cAMP. A representative ion chromatogram of cAMP showing the parent and daughter ion peaks for AtNCED3^211-440^ (see arrows). The inset shows an SDS-PAGE gel of the recombinant protein. (**D**) HPLC elution profile of cAMP generated by AtNCED3^211-440^. The calculated amount of cAMP after 25 minutes of enzymatic conversion of AtNCED3^211-440^ was 51 ± 4 fmol/μg protein (*n* = 3). The inset shows a calibration curve of the peak height plotted against fmoles of cAMP. (**E**) cAMP generated by AtNCED3^S311P/D313^ shows a strongly reduced peak as compared to the AtNCED3^211-440^. (**F**) Amount of cAMP produced after 25 minutes as a function of AtNCED3^211-440^ protein concentration. Values are means ± SD (n = 3).

For the *in vitro* assessment of AC activity, enzyme immunoassay (Fig. 2B) and mass spectrometry (LC-MS/MS) detection were performed. Both AtNCED3^211-440^ and the AtNCED3^S311P/D313T^ double mutant were expressed in *E. coli* and affinity purified (see Supplemental Methods). Their AC activities were tested in a reaction mixture containing ATP and Mn^2+^ as the cofactor. After 25 minutes of reaction, AtNCED3^211-440^ generated 45.4 ± 3 fmol cAMP/μg protein (8.809 × 10^−4^ fmol cAMP/pmol protein/s) while AtNCED3^S311P/D313T^ double mutant only yielded 4.3 ± 2 fmol cAMP/μg protein (7.328 × 10^−5^ fmol cAMP/pmol protein/s) which is approximately 10x lower; a difference that is statistically significant (*P*<0.005; *n*=3) (Fig. 2B). Measurements at each time point compared to the first time point for both AtNCED3^211-440^ and AtNCED3^S311P/D313T^ are also statistically significant (*P*<0.05; *n*=3).

Furthermore, cAMP levels were measured by LC–MS/MS specifically identifying the presence of the unique product ion at m/z 136 [M+H]^+^ that is fragmented in a second ionization step in addition to the parent ion at m/z 330 [M+H]^+^. This fragmented product ion was used for quantitation. A representative ion chromatogram of cAMP showing both the parent and product ion peaks is shown in Fig. 2C. AtNCED3^211-440^ generated 51 ± 4 fmol/μg protein of cAMP (*n*=3) after 25 minutes. This amount is not only consistent with the enzyme immunoassay measurements, but is also comparable with other experimentally characterized plant ACs that function as moonlighting proteins such as AtKUP7 (42.5 fmol/μg) [27] and AtKUP5 (38.1 fmol/μg) [28]. cAMP generated by the AtNCED3^S311P/D313^ also show a strongly reduced peak as compared to the AtNCED3^211-440^ in the LC-MS/MS detection (Fig. 2D, 2E). To further demonstrate the AC function of AtNCED3^211-440^, we conducted an independent set of experiments and show that the cAMP generation increases proportionally with the amount of enzyme (Fig. 2F and S2) recording a *V_max_* of 79.87 fmoles cAMP μg protein^−1^ and a *K_M_* of 0.863 mM, respectively (Fig. S3). Notably, the *K_m_* of NCED3 is comparable to the human soluble AC which is reported to be 0.8 mM although its *V_max_* is about one order of magnitude lower than that of the human soluble AC [30].

The seemingly low *in vitro* activities of these AC centers may be attributed to their moonlighting nature. Unlike canonical stand-alone enzymes, they only assume modulatory roles regulating the function of other primary domains in complex proteins to afford localized intrinsic regulation of protein domains within micro-environments of the plant cell such as by rapidly switching from one signaling pathway to another [31,32]. Furthermore, there is also a likelihood that components which are not present in the *in vitro* reaction mixture might significantly enhance enzymatic activity. One such component is calcium which we have found to significantly enhance NCED3 activity *in vitro* (Fig. S4). Thus, it is conceivable that cellular ion concentrations (of e.g., calcium and bicarbonate) could regulate the AC activity of AtNCED3 [30].

### 3.3 Meta-analysis of 9-cis-epoxycarotenoid dioxygenase 3

The dioxygenase activity of NCED3 requires the binding of Fe^2+^ octahedrally to four histidine residues (Fig. 1C) where ligated oxygen is used to cleave the aromatic rings of the 9-*cis*-epoxycarotenoid substrate to produce 2-*cis*,4-trans-xanthoxin and 12’-apo-carotenal, in the first step of ABA biosynthesis from carotenoids [33]. Through sequence analysis, the presence of corresponding histidines (Fig. 1A) would indicate if the Fe^2+^ binding and the AC catalytic centers co-locate and, in which case, implies that AtNCED3 has either dioxygenase or AC activity at any given time. Indeed, structural analysis of AtNCED3 and ZmNCED3 (PDB ID: 3NPE) showed high resemblance at their corresponding dioxygenase domains. The histidine residues of AtNCED3 are located deep into a pocket of the dioxygenase catalytic site much like in ZmNCED3 (Fig. 1C). Comparison of their ribbon models also revealed that the four iron coordinating histidines occupy similar spatial arrangements as in ZmNCED3. The surface models revealed that while the AC center is located at the solvent exposed region at the entrance of the dioxygenase pocket, there is however no physical obstruction or steric hindrance of the 9-*cis*-epoxycarotenoid substrate, with or without docking of ATP at the AC catalytic center (Fig. 1C).

The sequence and structural analysis imply that both activities could occur at the same time and could be independently regulated. We propose that AtNCED3 is a chloroplastic protein that acts at the intersection of two important signaling pathways involved in ABA and cAMP metabolisms, where both enzymatic activities have well-documented physiological effects including the regulation of plant osmosis, responses to salinity, energy metabolism and light stress responses [34–36]. Dual activity in moonlighting regulatory proteins is not uncommon especially in light of recent reports that have established catalytic centers of such nature (GCs and ACs) to perform a ‘tuning role’ in complex regulatory networks (e.g., [37,38]). While the biosynthesis and the molecular and biological functions of ABA are already well-understood, the same cannot be said for cAMP especially with respect to its function in the chloroplast. Experimental evidence as early as 1996 has confirmed that cAMP, but not cGMP and AMP, inhibits phosphorylation of proteins in the chloroplast especially those related to the light-harvesting chlorophyll a/b binding protein complex [39]. Later in 2004, cAMP was detected and quantified using a highly sensitive liquid chromatography/electrospray ionization tandem mass spectrometry method [40], which is also employed in this study. In 2005, the same group went on to demonstrate the topological AC activities in the chloroplast using cyto-enzymological and immune-cytochemical approaches. They have detected AC activities in the intermembrane space and importantly also in the stroma of the chloroplast [40] and since AtNCED3 is localized in the stroma, we may have identified the molecule that is responsible, at least in part, for the reported AC activities. Furthermore, the fact that AtNCED3 is partially bound to thylakoids [41] made this argument even more convincing because it would allow cAMP, which is known to operate in a transient and temporal manner within organelle micro-environments, to selectively exert the reported stronger phosphorylation inhibitory effects on proteins in the light-harvesting complex, that are also bound to the thylakoid membranes [39].

Taken together, we have shown experimental evidence for what could be the first AC reported in the chloroplast of higher plants and have assigned an important moonlighting role for AtNCED3. This adds to a growing list of plant proteins that are known to moonlight [24,29,31,42], where the role of some has already been characterized with direct connections to its primary domains thereby directly affecting e.g. ion transport [27,28] or enzyme activation or deactivation [14,32]. Our results will therefore guide future experimental work that focuses on the role of this AC in inter- and intra-molecular regulation in the chloroplast and, importantly, also its cellular and biological roles in higher plants. Future works can also employ this amino acid motif-based search strategy to identify candidate ACs in other species including *Homo sapiens* where structural analysis can then filter candidates for experimental studies *in vitro* and/or *in vivo*.

## Acknowledgments

We thank Salim Al-Babili (King Abdullah University of Science and Technology) for helpful discussion. We also thank Lee Staff for proofreading the manuscript.

