## Supplemental File for "Moonlighting adenylyl cyclases in plants – an *Arabidopsis thaliana* 9-*cis*-epoxycarotenoid dioxygenase as point in case"

**Supplemental Information contains four Figures, one Table, Supplemental Methods and Supplemental References.**

### Supplemental Figure S1

| **AtNCED3 ATP docking solutions** | | | | | | | |
|---|---|---|---|---|---|---|---|
| Mode 1st run | Affinity | Distance from best mode | | Mode 2nd run | Affinity | Distance from best mode | |
| | (kcal/mol) | rmsd l.b. | rmsd u.b. | | (kcal/mol) | rmsd l.b. | rmsd u.b. |
| 1 | -4.7 | 0.000 | 0.000 ✗ | 1 | -5.0 | 0.000 | 0.000 ✗ |
| 2 | -4.7 | 6.664 | 8.788 ✓ | 2 | -5.0 | 3.234 | 5.635 ✓ |
| 3 | -4.6 | 5.661 | 8.056 ✓ | 3 | -5.0 | 2.225 | 3.673 ✓ |
| 4 | -4.5 | 5.419 | 8.284 ✓ | 4 | -4.9 | 2.628 | 4.137 ✓ |
| 5 | -4.5 | 1.510 | 3.456 ✗ | 5 | -4.9 | 1.903 | 3.246 ✗ |
| 6 | -4.4 | 6.546 | 8.691 ✓ | 6 | -4.8 | 1.961 | 3.823 ✓ |
| 7 | -4.2 | 1.668 | 2.344 ✗ | 7 | -4.8 | 2.967 | 5.464 ✗ |
| 8 | -4.1 | 1.894 | 2.843 ✗ | 8 | -4.6 | 1.653 | 2.488 ✗ |
| 9 | -4.0 | 5.526 | 8.243 ✓ | 9 | -4.5 | 2.301 | 3.249 ✓ |
| Ave ± SD | -4.44 ± 0.27 | | | Ave ± SD | -4.84 ± 0.21 | | |
| **Mean ± SEM** | **-4.64 ± 0.03** | | | | | | |

✓ solutions with "correct binding pose" where adenine points towards S311 and phosphate towards K317
✗ solutions with "incorrect binding pose"

Only solutions with "correct binding pose" are considered in free energy calculations

**Total "correct binding pose" (✓) = 10/18 = 55.6 %**

**Correct ATP binding pose (10/18 ✓)** **Incorrect ATP binding pose (8/18 ✗)**

**1st run**

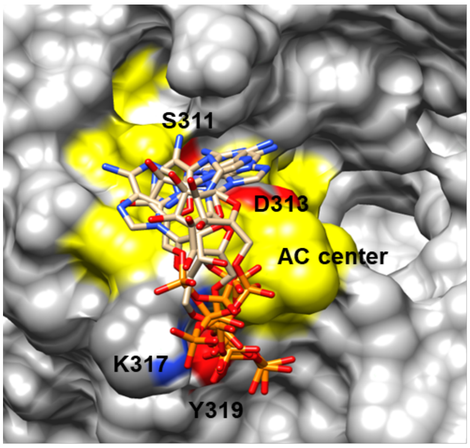

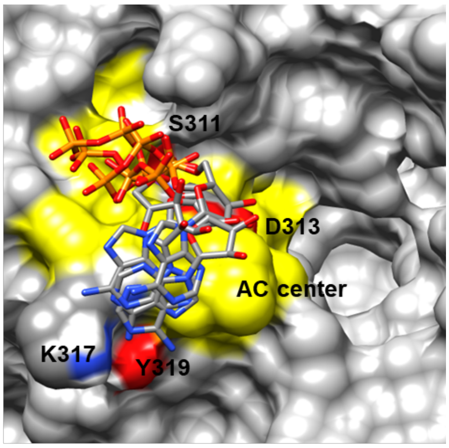

**2nd run**

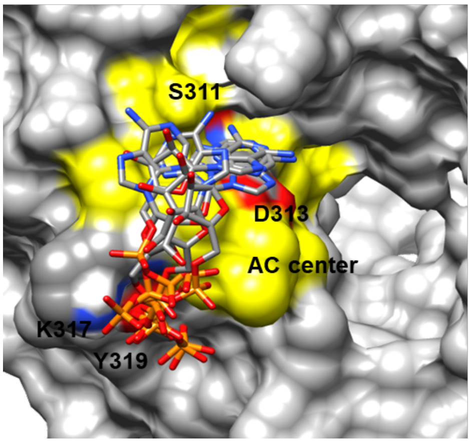

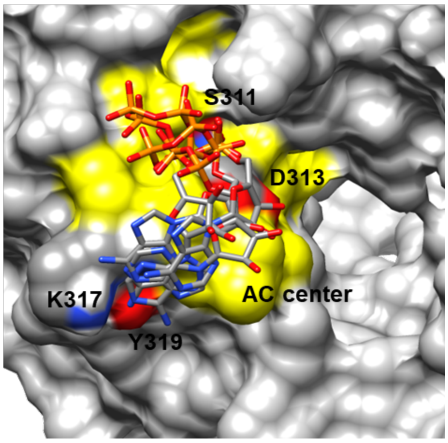

**AtNCED3 ATP docking clusters and data, and interpretation of the docking solutions.** A total of 18 solutions across two independent simulations generated by AutoDock Vina (ver. 1.1.2) [1] were evaluated in terms of free energies and binding poses. Orientations and binding poses were analyzed with the UCSF Chimera (ver. 1.10.1) [2]. Chimera is developed by the Resource for Biocomputing, Visualization, and Informatics at the University of California, San Francisco (supported by NIGMS P41-GM103311). In docking simulations, all bonds in the ATP ligand were allowed to move freely but the protein was set rigid. Docking orientations of ATP were evaluated based on a previously ascertained "correct binding pose" where the adenine of ATP points towards position 1 which normally occupies the interior of the AC pocket and, the phosphate which points towards position 14 which normally occupies the solvent-exposed entrance area of the AC pocket, respectively. Docking simulations consider both spatial and charge at the vicinity of the catalytic center based on pre-determined grids that cover the entire AC center and can afford free rotation of ATP substrate which we have set prior to docking experiments. Although varying ATP binding poses may have good binding affinities, not all docking solutions possess the "correct binding pose" that we have ascertained. Thus, we manually analyzed the binding pose of each solution and mark those with the correct binding poses as ✓ and those with incorrect binding poses as ✕. We also consider how frequent the software finds the “correct binding pose” and found that it is 55.6% with a mean binding affinity of -4.64 ± 0.03 kcal/mol.

**Supplemental Figure S2**

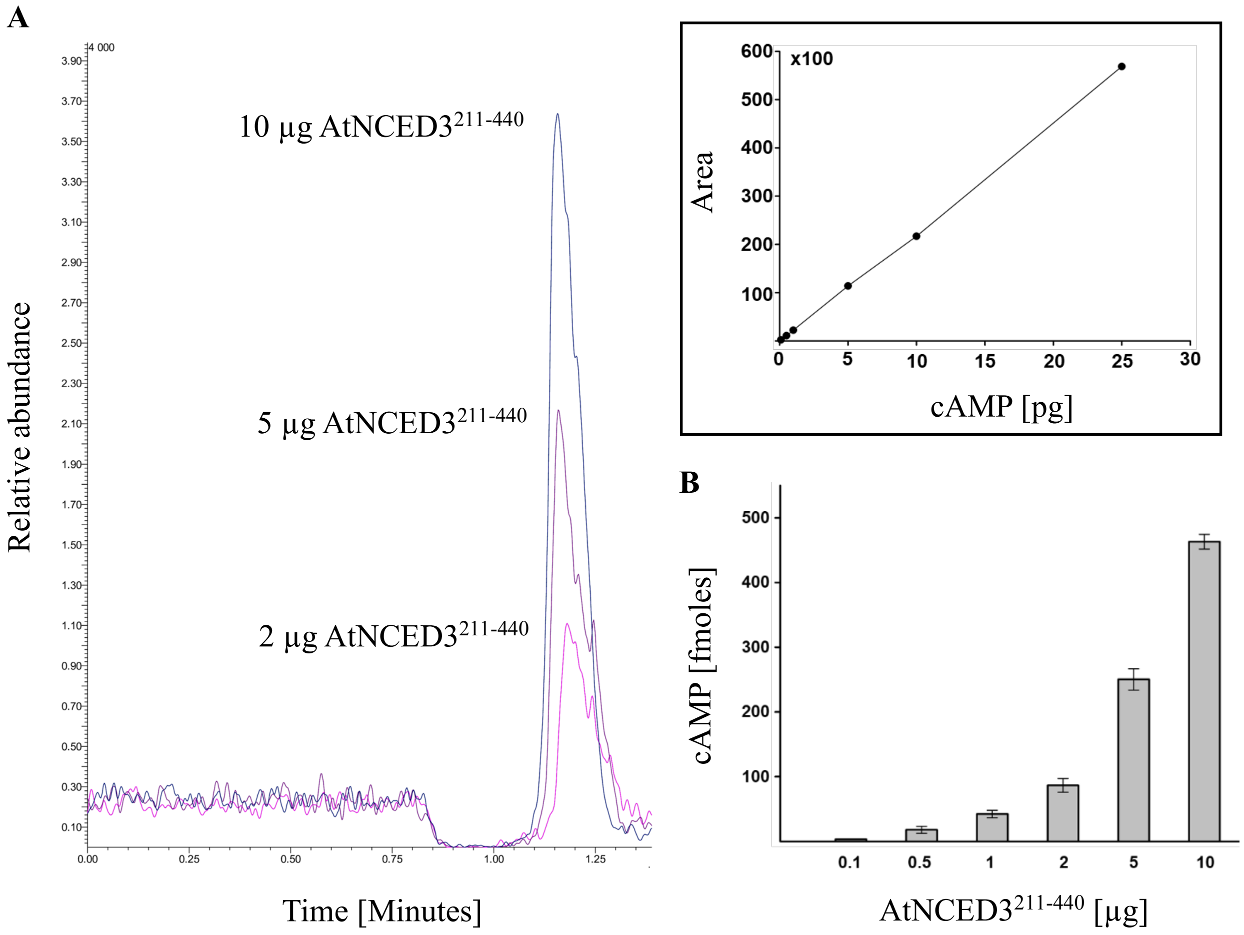

**Enzyme activity as a function of protein concentration.** (**A**) Representative MRM chromatograms of cAMP produced by AtNCED3$^{211\text{-}440}$ protein at concentrations of 2, 5 and 10 µg in the presence of 50 mM Tris-HCl pH 8, 2 mM IBMX, 1 mM ATP and 5 mM $MnCl_2$. Inset box shows the calibration curve for cAMP. LC-MS/MS experiments were performed using the Nexera UHPLC and LCMS-8045 integrated system (Shimadzu Corporation). The ionization source parameters were optimized in positive ESI mode using pure cAMP dissolved in HPLC-grade water (Sigma). Samples were separated using Ascentis® Express C18 HPLC Column (100 x 2.1 mm, 2.7 µm). An isocratic mix of solvent A (0.1 % (v/v) formic acid) and solvent B (100 % (v/v) methanol) (ratio: 90/10) was applied over 6 minutes with a flow rate of 0.3 mL/minute. The interface voltage was set at 4.0 kV for positive (ES+) electrospray. Data acquisition and analysis were done with LabSolutions workstation for LCMS-8045. (**B**) Amount of cAMP produced as a function of AtNCED3$^{211\text{-}440}$ protein concentration. Values are means ± SD ($n$ = 3).

### Supplemental Figure S3

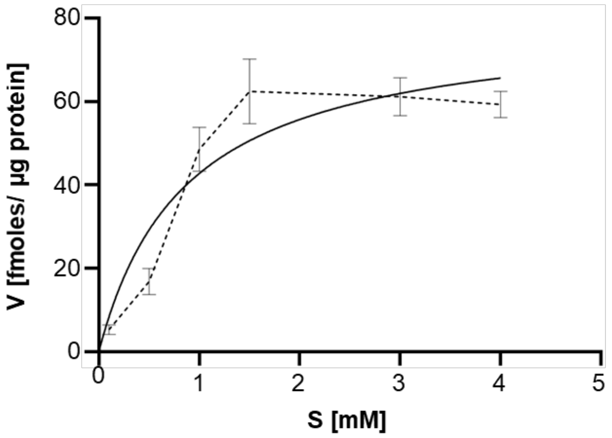

| | |
|---|---|
| Michaelis-Menten | |
| Best-fit values | |
| Vmax | **79,87** |
| Km | **0,8683** |
| 95% CI (profile likelihood) | |
| Vmax | 51,22 to 158,7 |
| Km | 0,1712 to 4,200 |
| Goodness of Fit | |
| Degrees of Freedom | 4 |
| R squared | 0,8788 |
| Sum of Squares | 376,4 |
| Sy.x | 9,700 |
| Constraints | |
| Km | Km > 0 |
| Number of points | |
| # of X values | 6 |
| # Y values analyzed | 6 |

**Calculation of the $K_M$ and $V_{max}$ of AtNCED3$^{211\text{-}440}$ *in vitro*.** Michaelis-Menten plot for adenylate cyclase activity of AtNCED3$^{211\text{-}440}$. The $V_{max}$ was 79.87 fmoles $min^{-1}$ µg $protein^{-1}$ and the $K_M$ was 0.8693 mM, respectively. Values are means ± SD ($n = 3$).

**Supplemental Figure S4**

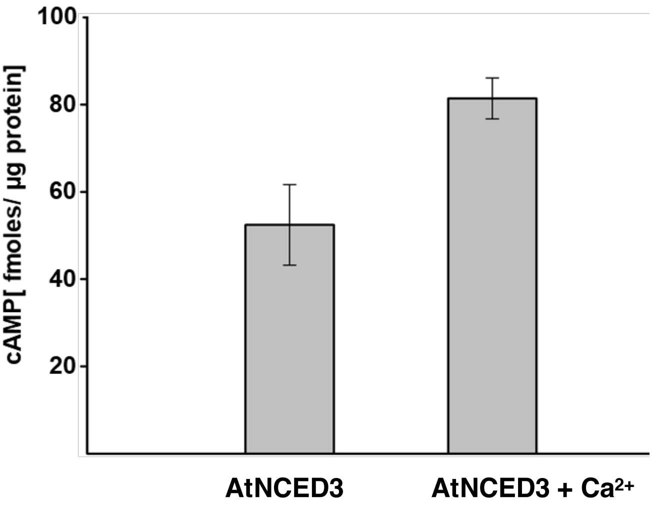

**Effect of $Ca^{2+}$ on AC activity of $AtNCED3^{211\text{-}440}$**. Cyclic AMP generated *in vitro* by 5 µg recombinant $AtNCED3^{211\text{-}440}$ protein in 25 minutes in the presence of 1 µM $Ca^{2+}$, 1 mM ATP, 5 mM $MnCl_2$, 2 mM IBMX and 50 mM Tris-HCl (pH 8). Data represent means ± SD ($n$ = 3).

### Supplemental Table

**Supplemental Table S1.** *Arabidopsis thaliana* proteins that contain the AC search term [RKS][YFW][DE][VIL]X{4}[Y]X{4}[KR]X{1,3}[DE].

| TAIR ID | Annotation |
|---|---|
| At1g30100 [1] | 9-*cis*-epoxycarotenoid dioxygenase, biosynth. of ABA - NCED5 |
| At1g47900 | Filament-like protein (DUF869) |
| At1g67120 | Homolog of the yeast MDN gene |
| At1g68110 [3] | CLAP, clathrin assembly protein, eap1 |
| At1g78390 [1] | 9-*cis*-epoxycarotenoid dioxygenase - NCED9 |
| At2g22560 | Kinase interacting (KIP1-like) family protein |
| At2g34520 [1] | Mitochondrial ribosomal protein S14, RPS14 |
| At3g14440 [1,2] | 9-*cis*-epoxycarotenoid dioxygenase - NCED3 |
| At4g18350 [1] | 9-*cis*-epoxycarotenoid dioxygenase - NCED2 |
| At5g59900 | Pentatricopeptide repeat (PPR) superfamily protein |
| At5g65160 | 36 carboxylate clamp (CC)-tetratricopeptide repeat (TPR) prot. |
| At5g67360 | Subtilisin-like serine prot. for mucilage release from seed coat |

[1] Proteins with a function in the biosynthetic process (GO:0009058); [2] Annotated as having a role in the hyperosmotic salinity response (GO:0042538) and the response to water deprivation (GO:0009414); [3] Experimentally confirmed AC.

### Supplemental Methods

#### Generation of recombinant AtNCED3$^{211\text{-}440}$

RNA was extracted from *A. thaliana* Col-0 leaf tissue using the RNeasy kit (Qiagen, Crawley, UK) and converted to cDNA using Superscript III Reverse Transcriptase according to the manufacturer's instructions (Invitrogen, Carlsbad, CA, US). Primers designed to amplify the AC domain of **AtNCED3$^{211\text{-}440}$** (forward: 5′-ATGATAGTCGACCCGGCACA-3′ and **AtNCED3$^{211\text{-}440}$**, reverse: 5′-TTAAGCATCAATCCACTTAATGTTCGA-3′). The cDNA was used as template in a PCR reaction with the **AtNCED3$^{211\text{-}440}$** AC primers and KAPA HiFi Taq Polymerase according to the manufacturer's instructions (KAPA Biosystems, Wilmington, MA, US). Subsequently, A overhangs were added using KAPA Taq Polymerase according to the manufacturer's instructions (KAPA Biosystems, Wilmington, MA, US) and the PCR product was cloned into the Gateway compatible pCR8 vector (Invitrogen, Carlsbad, CA, US). The AtNCED3$^{S311P/D313T}$ double mutant was generated by site directed mutagenesis using the following primers: AtNCED3$^{S311P/D318T}$ forward (5′- TCGCTTTAGGCTACTACGTCGTT-3′) and AtNCED3$^{S311P/D318T}$ reverse (5'-AACGACGTAGTAGCCTAAAGCGA-3′).

The AC domain of AtNCED3, and the double mutant were recombined into the pDEST17 expression vector (Invitrogen, Carlsbad, CA, US) to create pDEST17-AtNCED3$^{211\text{-}440}$ fusion constructs containing C-terminal His tags for affinity purification. These constructs were then transformed into *E. coli* cyaA mutants for functional complementation or *E. coli* BL21 A1 cells (Invitrogen, Carlsbad, CA, US) for recombinant protein expression. Purification of the recombinant proteins was performed under denaturing conditions using Ni-NTA agarose beads according to the manufacturer's instructions (Qiagen, Hilden, Germany) and refolded by Fast Protein Liquid Chromatography (FPLC) using HisTrap HP Ni-NTA columns (GE Healthcare, Little Chalfont, UK) as detailed in the next section.

#### Purification and refolding of recombinant AtNCED3

The recombinant cDNA encoding *AtNCED3$^{211\text{-}440}$* in the appropriate pDEST17-AtNCED3 $^{211\text{-}440}$ fusion construct was transformed into BL21 A1 *E. coli* cells (Invitrogen, Carlsbad, US) and grown in LB broth media containing 100 µg/mL ampicillin on an orbital shaker (New Brunswick Scientific, New Jersey, USA) at 200 rpm at 37°C, until the optical density ($OD_{600}$) reached 0.6. Recombinant protein expression was induced by adding 0.2 % l-arabinose and the culture grown for a further 4 hours at 37°C. The recombinant protein was purified by preparing a cleared cell lysate under denaturing conditions essentially as described in Protocols 10 and 17 of the QIAexpressionist manual (Qiagen, Crawley, UK) but with some modifications. Firstly, a cleared cell lysate was prepared by resuspending the harvested cells in lysis buffer (100 mM $NaH_2PO_4$, 10 mM Tris-Cl, 6 M guanidine hydrochloride; pH 8) at a ratio of 1 g pellet weight to 10 mL buffer volume and mixed with on a rotary mixer for 30 min and then centrifuged at 2300 x *g* for 15 minutes at room temperature. The cleared cell lysate supernatant was collected and mixed with 1 mL 50% (w/v) Ni-NTA slurry (Qiagen, Crawley, UK) that had been pre-equilibrated with 10 mL of lysis buffer. The contents were gently mixed on a rotary mixer (Breda Scientific, Breda, Netherlands) for 30 minutes at room temperature. The lysate-resin mixture was loaded into an empty PD-10 column (Amersham Pharmacia Biotech, Little Chalfont, UK), allowed to settle and the flow through discarded. The protein bound resin was washed three times with 30 mL wash buffer (8 M urea, 100 mM $NaH_2PO_4$, 10 mM Tris-HCl; pH 6.3) then with 2 mL elution buffer (8 M urea, 100 mM $NaH_2PO_4$, 10 mM Tris-HCl; pH 5.9) and fractions collected. The recombinant protein was subjected to a second elution with 2 mL imidazole-containing elution buffer (8 M urea, 100 mM $NaH_2PO_4$, 250 mM imidazole, 10 mM Tris-HCl; pH 8). The elution fractions that contained protein were pooled then desalted and concentrated to approximately 0.5 mL using the Amicon Ultra 15 Centrifugal Filter Unit, 15 kDa NMWL according to the manufacturer's instructions (Merck Millipore, Burlington, MA). This was diluted with 15 mL binding buffer (8 M urea, 20 mM $Na_2H_2PO_4$, 500 mM NaCl, 100 mM sucrose, 100 mM non-detergent sulfobetaines, 0.05% polyethylene glycol, 4 mM reduced glutathione, 0.04 mM oxidized glutathione and SIGMAFAST protease inhibitor cocktail at pH 7.8). The 1 mL HisTrap HP Ni-NTA column (GE Healthcare, Little Chalfont, UK) was connected to the AKTA Fast Protein Liquid Chromatography (FPLC) (GE Healthcare, Little Chalfont, UK) and equilibrated with 10 mL binding buffer at a flow rate of 1 mL/ minute. The denatured protein was then loaded on to the column at a flow rate of 0.2 mL/ minute. Once bound, the denatured protein was refolded by a gradual linear dilution of the 8 M urea to 0 M urea in refolding buffer (20 mM $Na_2H_2PO_4$, 500 mM NaCl, 500 mM sucrose, 100 mM non-detergent sulfobetaines, 0.05% PEG, 4 mM reduced glutathione, 0.04 mM oxidized glutathione and SIGMAFAST protease inhibitor cocktail at pH 7.8) at a flow rate of 1 mL/ minute for 50 column volumes. After renaturation, the column was washed with 10 column volumes of refolding buffer. Finally, the protein was eluted in a linear gradient for 20 column volumes with elution buffer (20 mM $Na_2H_2PO_4$, 500 mM NaCl, 500 mM sucrose, 500 mM imidazole, 100 mM NDSB, 0.05% PEG, 4 mM reduced glutathione, 0.04 mM oxidized glutathione and SIGMAFAST protease inhibitor cocktail at pH 7.8). Fractions containing the recombinant protein were pooled then de-salted and concentrated using the Amicon Ultra 15

Centrifugal Filter Unit, 15 kDa NMWL according to the manufacturer's instructions (Merck Millipore, Burlington, MA). The protein concentration was determined by the Bradford method [3] and the recombinant protein was stored at -20°C.

**Statistical analysis**

Statistical analysis was performed using an unpaired, one-tailed Student's *t*-test (Two-Sample Assuming Unequal Variances). Measurements are represented as mean ± SE and significance were set to a threshold of $P < 0.05$ and $n$ values represent number of experimental replicates. Measurements of AtNCED3$^{211\text{-}440}$ and AtNCED3$^{S311P/D313T}$ at each time point are compared to that of the first time point, and measurements of AtNCED3$^{S311P/D313T}$ double mutant is also compared to that of AtNCED3$^{211\text{-}440}$ at the corresponding time points.
